# Two novel 7-*epi*-zingiberene derivatives with biological activity from *Solanum habrochaites* are produced by a single cytochrome P450 monooxygenase

**DOI:** 10.1101/2020.04.21.052571

**Authors:** Sebastian Zabel, Wolfgang Brandt, Andrea Porzel, Benedikt Athmer, Ruy Kortbeek, Petra Bleeker, Alain Tissier

## Abstract

Secretions from glandular trichomes potentially protect the plant against a variety of aggressors. In the tomato genus, wild species constitute a rich source of chemical diversity produced at the leaf surface by glandular trichomes. Previously, 7-*epi*-zingiberene produced in several accessions of *Solanum habrochaites* was found to confer resistance to whiteflies (*Bemisia tabaci*) and other insect pests. Here, we identify two derivatives of 7-*epi*-zingiberene from *S. habrochaites* that had not been reported as yet. We identified them as 9-hydroxy-zingiberene and 9-hydroxy-10,11-epoxyzingiberene. Using a combination of genetics and transcriptomics we identified a single cytochrome P450 oxygenase, ShCYP71D184 that carries out two successive oxidations to generate the two sesquiterpenoids. Bioactivity assays showed that only 9-hydroxy-10,11-epoxyzingiberene exhibits substantial toxicity against *B. tabaci*. In addition, both 9-hydroxy-zingiberene and 9-hydroxy-10,11-epoxyzingiberene display substantial growth inhibitory activities against a range of microorganisms, including *Bacillus subtilis*, *Phytophtora infestans* and *Botrytis cinerea*. Our work shows that trichome secretions from wild tomato species can provide protection against a wide variety of organisms. In addition, the availability of the genes encoding the enzymes for the pathway of 7-*epi*-zingiberene derivatives makes it possible to introduce this trait in cultivated tomato by precision breeding.

## Introduction

The plant surface constitutes the first barrier against aggressions of various kinds, including pathogens and herbivores. In the aerial parts, different kinds of protuberances at the surface differentiate and contribute to the protection of plants against different types of stresses. The most prominent of these are trichomes, which are uni- or multicellular structures derived from the epidermis that can either be glandular or non-secreting. Glandular trichomes typically contain one to several highly metabolically active cells dedicated to the production of large quantities of secondary metabolites. The compounds produced can be either stored in a specialized cavity, which is typical for volatiles, or secreted onto the leaf surface as in the case of the resinous acyl sugars or diterpenoids in several Solanaceae species. In the tomato genus, including in the cultivated and related wild species, different types of glandular trichomes specialized in the production of distinct classes of substances can be found. Type I/IV trichomes typically produce acyl sugars, which are directly secreted from the tip of the glandular cells (King et al., 1990; Schilmiller et al., 2012). Type VI glandular trichomes constitute one of the most abundant types and in cultivated tomato, *Solanum lycopersicum*, their main products are monoterpenes (Schilmiller et al., 2009). Whereas little, if any, variation in the composition of the type VI secretion has been reported within *S. lycopersicum accessions*, related species, in particular *Solanum habrochaites*, display an impressive diversity of chemotypes (Gonzales-Vigil et al., 2012). Except in some accessions of *S. habrochaites ssp. glabratum*, where the main compounds produced are methyl ketones, which are fatty acid derivatives (Farrar and Kennedy, 1987; Fridman et al., 2005), monoterpenoids and sesquiterpenoids are the major substances produced by type VI trichomes. Based on the nature of the farnesyl diphosphate (FDP) precursor two main classes of sesquiterpenoids have been described. Class I sesquiterpenoids are synthesized from the cytosolic *trans,trans*-FDP (*E,E*-FDP) and consist of a mixture of various germacrene sesquiterpenes, as well as farnesoic and/or dehydro-farnesoic acids in some accessions (Snyder et al., 1993; Breeden et al., 1996; van Der Hoeven et al., 2000). Class II sesquiterpenes are derived from the *cis,cis*-FDP precursor (*Z,Z*-FDP) and are produced in the plastids (Sallaud et al., 2009). Until now two sesquiterpene synthases able to convert *Z,Z-*FPP have been identified in *S. habrochaites*. The santalene and bergamotene synthase (ShSBS) is a multiproduct cyclase which makes (+)-α-santalene, (+)-*endo*-β-bergamotene, and (-)-*endo*-β-bergamotene as well as other minor products (Sallaud et al., 2009). The other sesquiterpene cyclase (ShZS) synthesizes 7-*epi*-zingiberene (Figure 1, peak **1**) as the main product, which can spontaneously oxidize to *ar*-curcumene (Bleeker et al., 2012; Gonzales-Vigil et al., 2012). Accessions that make santalene and bergamotene do not produce 7-*epi*-zingiberene and *vice versa*, indicating that the corresponding enzymes are encoded by alleles of the same gene (Gonzales-Vigil et al., 2012). The santalene and bergamotene sesquiterpenes are further oxidized to carboxylic acids, which are, in some accessions such as LA1777, by far the most abundant compounds produced in type VI trichomes, with amounts that can reach tens of mg per leaf fresh weight (Frelichowski and Juvik, 2005). For all these compounds, activities towards various insect herbivores could be demonstrated. For example, the sesquiterpene carboxylic acids from LA1777 were shown to confer resistance against two lepidopteran pests, *Helicoverpa zea* and *Spodoptera exigua* (Frelichowski and Juvik, 2001). 7-*epi*-zingiberene seems to confer a particularly broad range of resistance against various pests, including pinworms (*Tuta absoluta*), whiteflies (*Bemisia tabaci*), spider mite (*Tetranychus evansi*), and Colorado potato beetle (*Leptinotarsa decemlineata*) (Carter et al., 1989; Maluf et al., 2001; Freitas et al., 2002; de Azevedo et al., 2003; Goncalves et al., 2006; Maluf et al., 2010). Transformation of cultivated tomato (*S. lycopersicum*) with the two genes required for the biosynthesis of 7-*epi*-zingiberene also resulted in improved insect-resistance due to an increased repellence of whiteflies and toxicity to spidermites (Bleeker et al., 2012).

**Figure 1.**
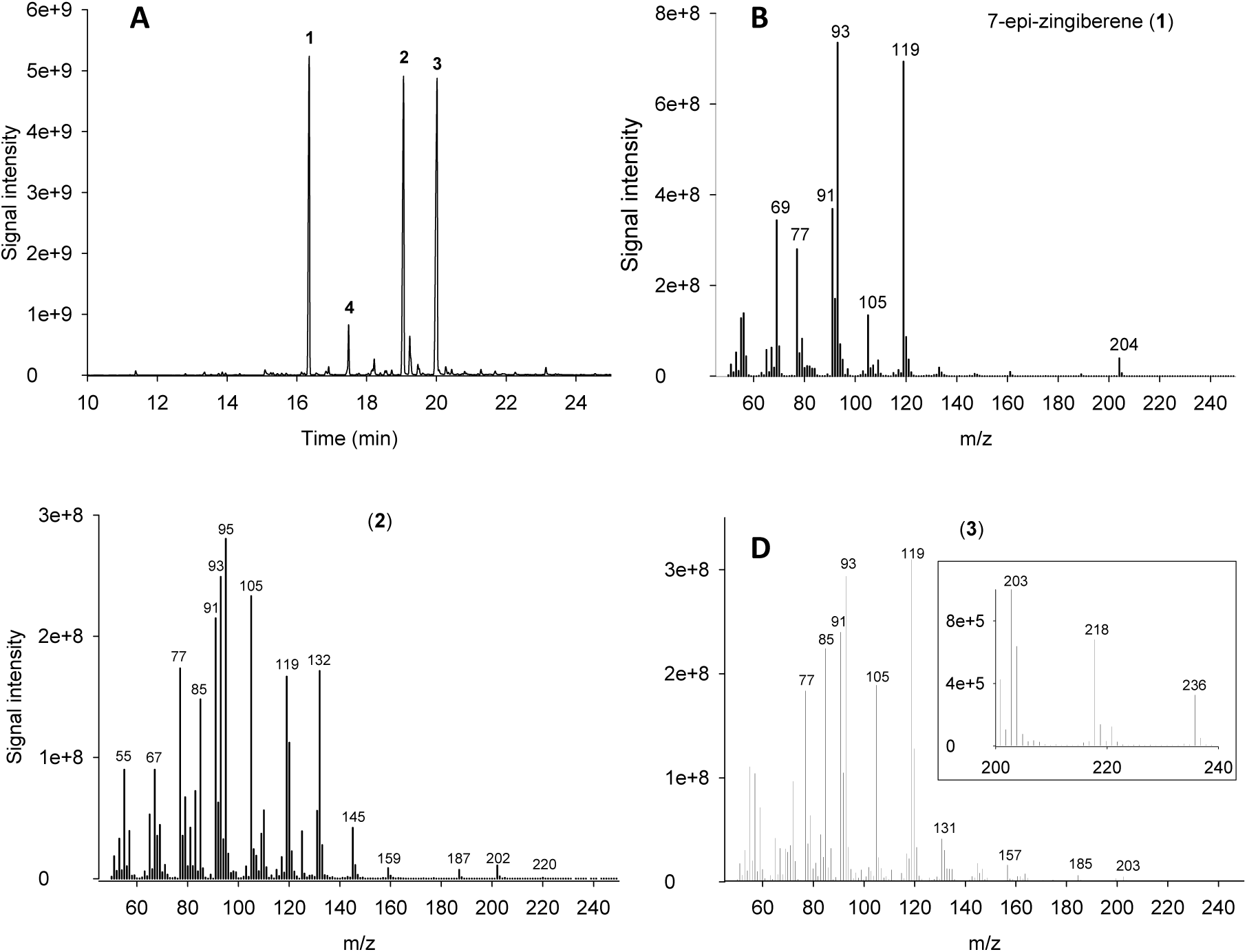
GC-MS analysis of leaf surface extracts of Solanum habrohaites LA2167. (A) GC-MS total ion chromatogram (TIC) of a leaf surface hexane extract of S. habrochaites LA2167. Peak 1 is 7-epi-zingiberene, peaks 2 and 3 are zingiberene derivatives which are characterized in this work, peak 4 is germacrene B. (B) Mass spectrum of 7-epi-zingiberene. (C) Mass spectrum of 2. (D) Mass spectrum of 3. The insert is a zoomed-in section between m/z 200 and 240.

While surveying the exudate profiles of various accessions of *S. habrochaites*, we noted the presence of additional peaks in accession LA2167 that were suggested to be derivatives of 7-*epi*-zingiberene based on their mass fragmentation pattern. We describe the structure elucidation of these compounds based on a combination of mass spectrometry, nuclear magnetic resonance spectroscopy (NMR) and circular dichroism. The derivatives were identified as (4*R*,7*R*,9*S*)-7-*epi*-9-hydoxy-zingiberene and (4*R*,7*R*,9*S,10S*)-*7*-*epi*-9-hydroxy-10,11-epoxy-zingiberene. Furthermore, using genetic and transcriptomics approaches we could identify a single gene coding for a cytochrome P450 oxygenase that was responsible for their biosynthesis. Finally, bioassays for antibacterial, fungicide and insecticide activities showed that these compounds display a range of biological activities, supporting their role in the defense of tomato plants against various pests.

## Results

### Identification and structure elucidation of two novel 7-*epi*-zingiberene derivatives

A metabolic profiling survey of different accessions from the wild tomato *Solanum habrochaites* revealed two novel major peaks in leaf surface extracts of LA2167 measured by GC-MS (Figure 1). The mass spectra of these compounds did not yield any convincing match in the NIST database, indicating that these might be novel compounds. However, the spectra did present significant similarities to that of *7-epi*-zingiberene, suggesting they might be derivatives thereof, with a molecular mass of 220 for compound **2** and 236 for compound **3** (Figure 1). This increase in molecular mass corresponds to one and two oxygen atoms for **2** and **3**, respectively, and is consistent with increased retention times upon gas chromatography. To further characterize these compounds, leaf surface extracts were derivatized by silylation and analyzed by GC-MS. Derivatization of **2** afforded a mass spectrum with a molecular ion of 292, a base peak of 197 and distinct m/z signals of 277 and 157 (**Supplemental Figure 1A**). These signals could be accounted for by the presence of a single hydroxyl group at C9 as seen in **Supplemental Figure 2**. Silylation of **3** increased the molecular mass by only 72, indicating that only one silylation had occurred (**Supplemental Figure 1B**). Since **3** has two additional oxygen atoms, this suggests that one of them is not available for derivatization and therefore could be, for example, present as an epoxide. One interpretation of the fragmentation pattern suggests the presence of a hydroxyl group at position 9, as in **2**, and of an epoxide between C10 and C11, instead of the double bond present in zingiberene for **3** (**Supplemental Figure 3**).

We purified **2** and **3** they from hexane extracts made from approximately 1000 leaves of *S. habrochaites* LA2167 in a single step on an open 50 mL silica column (see methods section and **Supplemental Figure 4**). After purification, 21 mg of **2** and 25 mg of **3** were recovered. **2** and **3** were analyzed by electrospray high resolution mass spectrometry (ESI-HRMS) in the positive mode giving monoisotopic molecular masses of 221.1900 and 237.1849 respectively, confirming the predicted formula C_15_H_24_O for **2** and C_15_H_24_O_2_ for **3** (**Supplemental Figures 5 and 6, Supplemental Tables 1 and 2**).

The structures of **2** and **3** were elucidated on the basis of extensive 1D (^1^H, ^13^C) and 2D (COSY, HSQC, HMBC,) NMR spectroscopic analysis. The ^13^C and DEPT spectra (Table 1) of **2** exhibited signals for four methyl, two methylene and seven methine groups as well as two quaternary carbons. As can be seen from their low-field ^13^C chemical shifts, the quaternary (δ ^13^C 133.9, 131.2 ppm) and four of the tertiary carbons (δ ^13^C 129.8, 129.6, 128.8, 120.9 ppm) belong to double bond units. One of the two remaining methine groups shows a hydroxyl substituent (δ ^13^C 67.0 ppm; δ ^1^H 4.356 ppm). Detailed analysis of ^1^H,^1^H COSY and ^1^H,^13^C HMBC 2D spectra (**Supplemental Figure 7** and Table 1) and comparison with literature data (Ishii et al., 2011) revealed **2** to be 9-hydroxy-7-epi-zingiberene, which we henceforth refer to as 9HZ (Figure 2).

**Figure 2.**
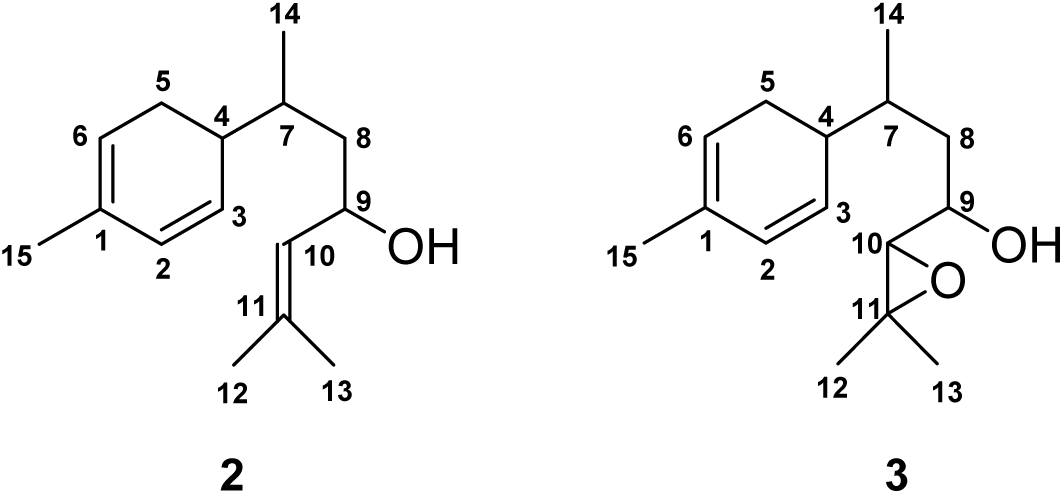
Structure of compounds 2 and 3 as determined by NMR spectroscopy. **2** and **3** were determined to be 9-hydroxy-zingiberene (9HZ) and 9-hydroxy-10,11-epoxy-zingiberene respectively.

**Table 1.** NMR data of Compound 2 (solvent: C_6_D_6_; numbering scheme see Figure 2)

| Pos. | $\delta^{13}\text{C}$ [ppm] | $\delta^1\text{H}$ [ppm] m (J [Hz]) | HMBC correlation<br>H to C |
| --- | --- | --- | --- |
| 1 | 131.2 | --- |  |
| 2 | 128.8 | 5.829 ddd (9.7/3.4/1.7) | 3, 4, 6 |
| 3 | 129.6 | 5.684 dd (9.7/3.4) | 1, 2, 4, 5, 7 |
| 4 | 38.7 | 2.377 m |  |
| 5 | 26.5 | 2.04<sup>a</sup> | 1, 3, 4, 6, 7 |
| 6 | 120.9 | 5.427 m | 15 |
| 7 | 33.5 | 1.66<sup>a</sup> | 3, 4, 5, 8, 14 |
| 8 | 42.3 | 1.553 ddd (13.4/7.2/5.2);<br>1.496 ddd (13.4/8.6/6.7) | 4, 7, 9, 10, 14 |
| 9 | 67.0 | 4.356 ddd (8.7/7.2/6.7) | 11 |
| 10 | 129.8 | 5.110 d sept(8.7/1.4) | 12, 13 |
| 11 | 133.9 | --- |  |
| 12 | 25.7 | 1.557 d (1.4) | 10, 11, 13 |
| 13 | 18.0 | 1.475 d (1.4) | 10, 11, 12 |
| 14 | 17.6 | 0.910 d (6.8) | 4, 7, 8 |
| 15 | 21.2 | 1.663 br s | 1, 2, 6 |

<sup>a</sup> $^1\text{H}$ chemical shift extracted from the HSQC spectrum

The molecular formula of **3** established by ESI-HRMS is C_15_H_24_0_2_, indicating one additional oxygen atom but the same number of four double bond equivalents as 9HZ. The NMR spectra of **3** largely resemble those of 9HZ (Table 2). However, the signals of CH-10 (δ ^13^C 128.9 ppm; δ ^1^H 5.110 ppm) and C-11 (δ ^13^C 133.9 ppm) in 9HZ are replaced in **3** by signals at δ ^13^C 68.1 ppm; δ ^1^H 2.558 ppm and δ ^13^C 58.9 ppm, respectively. The coupling constant of C-10/H-10 was extracted from the residual 1J correlation peak in the HMBC spectrum. Its size of 170 Hz with simultaneous high-field shift of H-10 clearly shows CH-10 to be involved in an epoxy unit. The structure of **3** was thus established as 9-hydroxy-10,11-epoxy-zingiberene, which we now refer to as 9H10epoZ (Figure 2). Notably, the NMR data was not sufficient to determine the absolute configuration of 9HZ and 9H10epoZ. This could be addressed after identification of the enzyme responsible for their synthesis (see below).

**Table 2.** NMR data of Compound 3 (solvent: C_6_D_6_; numbering scheme see Figure 2)

| Pos. | $\delta^{13}\text{C}$ [ppm] | $\delta^1\text{H}$ [ppm] m (J [Hz]) | HMBC correlation H to C |
| --- | --- | --- | --- |
| 1 | 131.2 | --- |  |
| 2 | 129.0 | 5.822 ddd (9.8/3.4/1.7) | 1, 4, 6, 15 |
| 3 | 129.0 | 5.618 dd (9.8/3.3) | 1, 4, 5, 6, 7 |
| 4 | 37.8 | 2.346 d | 3, 9 |
| 5 | 26.6 | 2.02<sup>a</sup> | 1, 3, 4, 6, 7 |
| 6 | 121.0 | 5.419 m | 4, 5, 15 |
| 7 | 32.9 | 1.748 m | 4, 5, 8, 9, 14 |
| 8 | 38.4 | 1.476 ddd (14.0/6.6/5.2);<br>1.435 ddd (14.0/8.3/7.0) | 4, 7, 9, 10, 14 |
| 9 | 69.0 | 3.442 ddd (8.3/7.8/5.2) | 7, 8, 10, 11 |
| 10 | 68.1 | 2.558 d (7.8) | 8, 9, 11, 12 |
| 11 | 58.9 | --- |  |
| 12<sup>b</sup> | 24.8 | 1.058 s | 10, 11, 13 |
| 13<sup>b</sup> | 19.4 | 1.046 s | 10, 11, 12 |
| 14 | 18.2 | 0.906 d (6.9) | 4, 7, 8 |
| 15 | 21.2 | 1.663 br s | 1, 2, 6 |

<sup>a</sup> $^1\text{H}$ chemical shift extracted from the HSQC spectrum; <sup>b</sup> may be interchanged

### Identification of a candidate gene for the biosynthesis of the oxidized zingiberene derivatives

To identify the genes involved in the biosynthesis of the two zingiberene derivatives, 9HZ and 9H10EPOZ, two complementary approaches were used: transcriptomics and genetics. For transcriptomics, RNA microarray hybridization was performed on different accessions of *S. habrochaites* that differ in their sesquiterpene composition. Among those that we selected, LA2167 is the only one that produces the oxidized zingiberenoids. We searched for genes annotated as oxidases, particularly cytochrome P450 oxygenases (CYPs) that are overexpressed in LA2167 compared to the other accessions. This analysis produced a set of three CYP-encoding genes, one of which showed a particularly strong expression in LA2167, *Sohab01g008670* (defined by similarity to the Solyc01g008670 gene of *S. lycopersicum*) (Figure 3). The expression of this gene was then verified by quantitative RT-PCR, which confirmed the particularly high expression of Sohab01g008670 in accession LA2167 (Figure 3).

**Figure 3.**
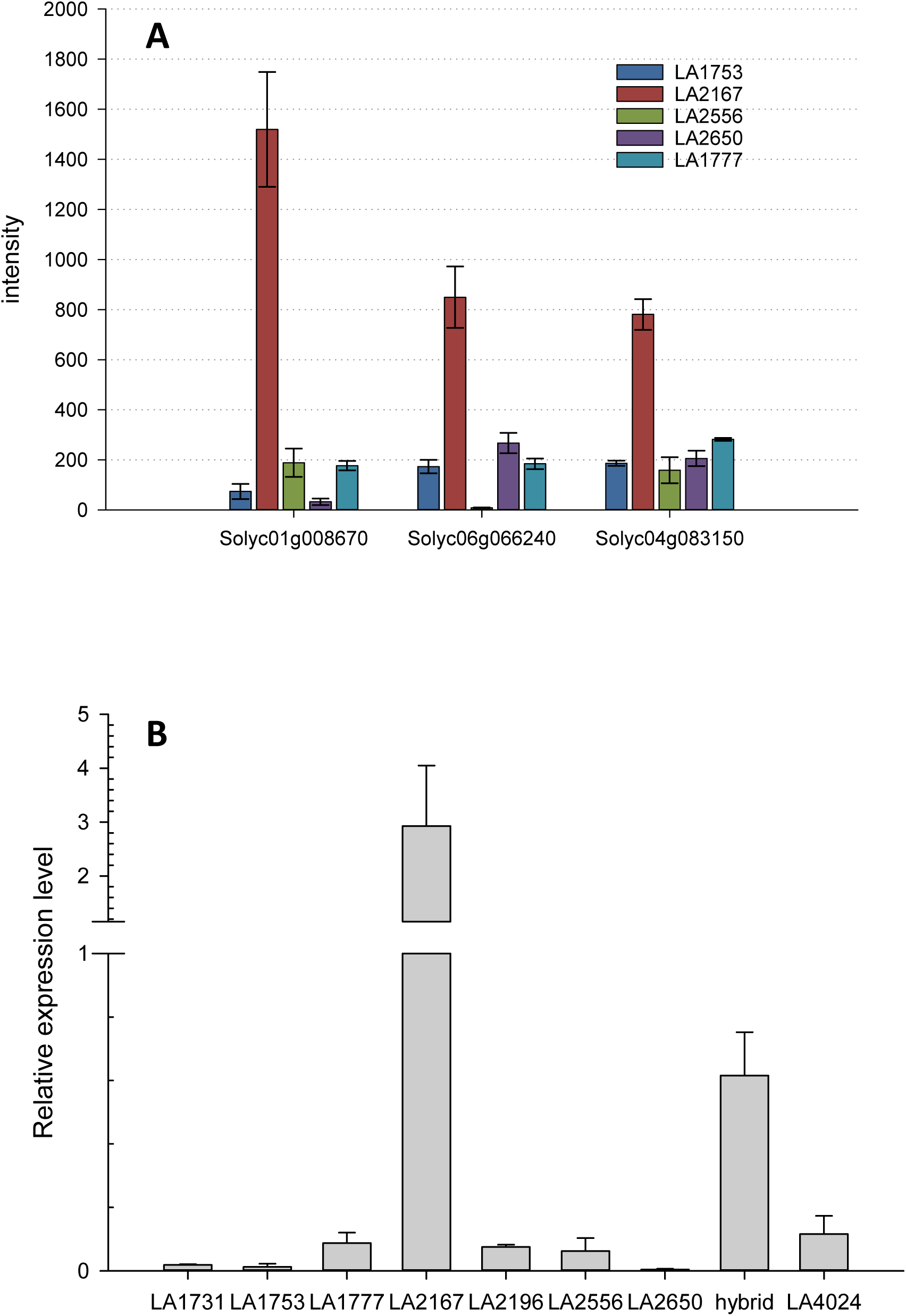
Expression of CYP-encoding candidate genes for the oxidation of zingiberene in various *S. habrochaites* accessions. **A**. Microarray data of 3 CYP encoding genes in 5 different accessions. LA2167 is the only accession that produces zingiberene and associated derivatives. Intensity designates the normalized fluorescence signal from the microarray **B**. qRT-PCR of Sohab01g008670 in different tomato accessions. Hybrid designates F1 plants of a cross between LA2167 *(S. habrochaites*) and LA4024 (*S. lycopersicum*).

In parallel, we generated a backcross population (BC1F1) (n=150) between *S. habrochaites* LA2167 and *S. lycopersicum* LA4024, using LA4024 as the recurrent parent. F1 plants exhibited an exudate profile that was a combination of that of both parents, including 9HZ and 9H10epoZ, indicating that the biosynthesis of the zingiberene derivatives is a dominant trait. Profiling of the exudate of the 150 BC1 plants by GC-MS showed a mendelian segregation for zingiberene and its derivatives (see Table 3). Notably, both zingiberene derivatives strictly co-segregated indicating that a single locus was responsible for the biosynthesis of these compounds. Furthermore, approximately half of the zingiberene producing lines also produced the zingiberene derivatives, demonstrating that these two loci are not linked. The BC1 population was genotyped using a set of 115 markers distributed over the 12 chromosomes. Linking the presence of 9HZ and 9H10EPOZ to genetic markers indicated that the locus for the biosynthesis of the zingiberene derivatives is on chromosome 1, in a region that includes *Sohab01g008670* (see **Supplemental Table 1** for the full set of data). Combined with the gene expression analysis, the genetic mapping data provided strong support for Sohab01g008670 as the gene responsible for the oxidation of zingiberene. According to the cytochrome P450 oxygenase nomenclature, Sohab01g008670 was named ShCYP71D184.

**Table 3.** Segregation of 7-*epi*-zingiberene, 9HZ and 9H10epoZ in the backcross population

| Metabolite | Number of backcross plants |  |  | Chi-Square test (n=1) |
| --- | --- | --- | --- | --- |
|  | 73 | 41 | 36 |  |
| 7-*epi*-zingiberene | - | + | + | 0.75 |
| 9HZ | - | - | + | 0.53 |
| 9H10epoZ | - | - | + |  |

### ShCYP71D184 is a zingiberene oxidase (ShZO)

To evaluate the enzymatic activity of ShCYP71D184, we expressed the full-length coding sequence transiently in *Nicotiana benthamiana* together with the enzymes required for the production of 7-*epi*-zingiberene (hereafter 7epiZ), namely the *Z,Z*-FPP synthase (zFPS) and the 7-*epi*-zingiberene synthase (ZS) from *S. habrochaites* (Sallaud et al., 2009; Bleeker et al., 2012). Leaf surface extracts of agro-infiltrated *N. benthamiana* leaves were analyzed by GC-MS five days after infiltration. We detected 7epiZ and small amounts of *ar*-curcumene in leaf surface extracts of *N. benthamiana* leaves expressing zFPS and ZS (Figure 4). When ShCYP71D184 was co-expressed with zFPS and ZS, several additional peaks appeared in the GC-MS chromatogram. Comparison with a leaf surface extract of LA2167 revealed that both 9HZ and 9H10epoZ were present (Figure 4 and **Supplemental Figure 8**). There was one additional major peak, whose abundance varied between repetitions, but it was typically higher than 9H10epoZ. This unidentified product is also present in LA2167 leaf surface extracts, although there it is less abundant than 9HZ and 9H10epoZ (Compound 6 in Figure 4). The similarity of the mass spectrum of the unidentified compound to that of *ar-*curcumene and its molecular mass of 234 (**Supplemental Figure 8**) suggest it could be 9-hydroxy-10,11-epoxy-curcumene. The absence of these zingiberene oxidation products in all other enzyme combinations or controls therefore confirms the function of ShCYP71D184 as a zingiberene oxidase, henceforth abbreviated as ShZO. A phylogenetic analysis of ShZO shows that it is most closely related to CYP71 enzymes from the Solanaceae, such as premnaspirodiene oxygenase (CYP71D55) and *epi*-aristolochene 1,3-dihydroxylase (CYP71D20) (**Supplemental Figure 9**). Notably, both CYP71D55 and CYP71D20 carry out successive oxidations of sesquiterpenes (Ralston et al., 2001; Takahashi et al., 2005; Takahashi et al., 2007).

**Figure 4.**
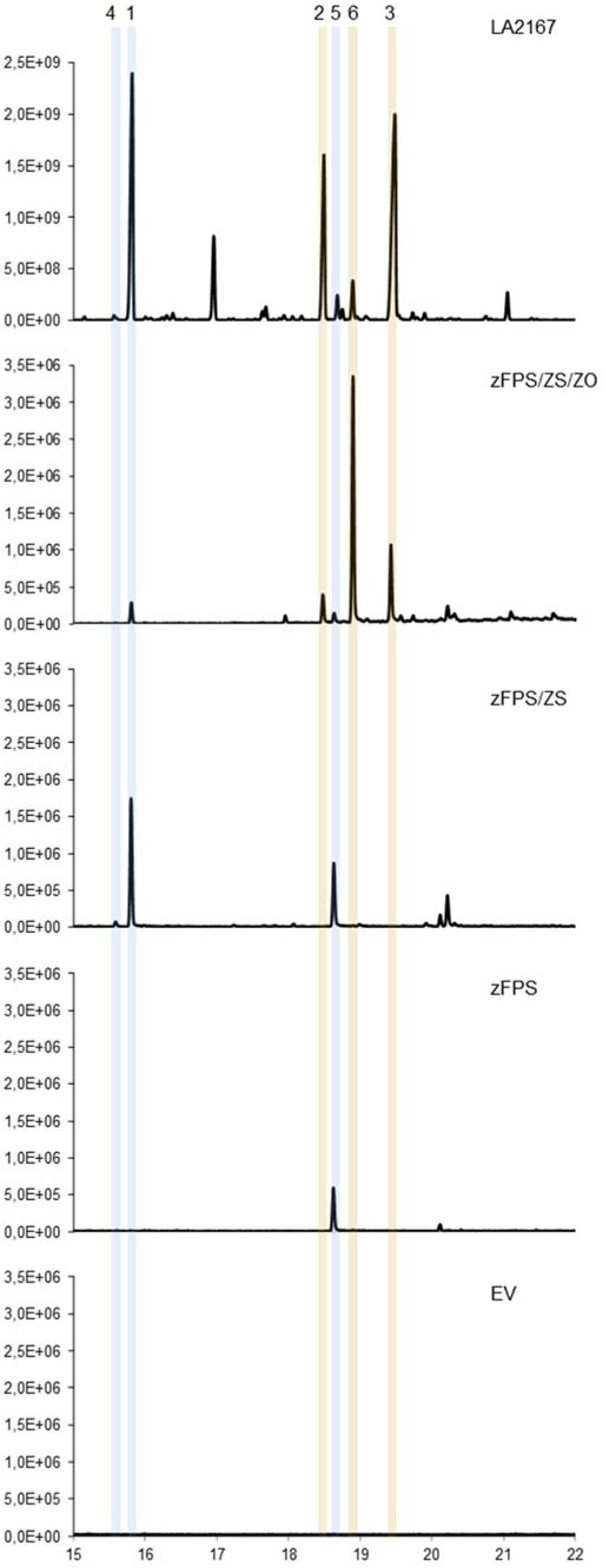
Expression of ShZS and ShCYP71D184 in *Nicotiana benthamiana*. The gene combinations indicated at the right of each chromatogram were agro-infiltrated in *N. benthamiana* and leaf extracts analysed by GC-MS. The portion of the chromatograms covering sesquiterpenes is shown. LA2167: leaf surface extracts of *S. habrochaites* LA2167. zFPS: *cis,cis*-farnesyl diphosphate synthase; ZS: zingiberene synthase; ZO: ShCYP71D184 (zingiberene oxidase). The detected compounds are labelled as follows. 1: 7-epizingiberene; 2: 9-hydroxy-zingiberene; 3: 9-hydroxy-10,11-epoxyzingibernee; 4: ar-curcumene; 5: farnesol; 6: unknown compound, tentatively identified as 9-hydroxy-10,11-epoxy-ar-curcumene

### Determination of the absolute configuration of 9HZ and 9H10epoZ by modelling

The NMR data did not allow us to determine the absolute configuration of 9HZ and 9H10epoZ. In the absence of crystals, which could have been used for X-ray spectroscopy, we decided to tackle this issue by two modelling approaches. The first consists in the comparison of simulated of circular dichroism (CD) spectra with the actual measured spectra, and the second involves substrate docking simulations on a modelled 3D structure of ShZO. Previous work established the absolute configuration of 7epiZ at carbon 4 as *R* (Bleeker et al., 2011). None of the calculated CD-spectra with the *S4* configuration fit the measured spectra, thus confirming the determination of configuration by Bleeker et al. (2011). Calculated CD-spectra for 9HZ with the configurations with *R*4, *R*7,*R*9 and *R*4, *S*7,*R*9 (**Supplemental Figures 10a** and **10c**) do not fit the experimental spectra and can therefore be rejected. The configuration with *R*4,*R*7,*S*9 (**Supplemental Figure 10b**) fits slightly better than the one with *R*4,*S*7,*S*9 (**Supplemental Figure 10d**). In the case of 9H10epoZ, out of all eight configurations only *R*4,*S*7,*R*9,*S*10 (**Supplemental Figure 11f)** can be excluded. This is in agreement with the poor fit for *R*4, *S*7,*R*9 of 9HZ (see above). All other configurations fit rather nicely with the respective experimental spectra (**Supplemental Figure 11**). Taking into account that 9H10epoZ is derived from 9HZ and that the simulated CD spectra of the *R*4,*R*7,*R*9 and *R*4,*S*7,*R*9 configurations do not fit, the configurations with *R*4,*R*7*,R*9,*S*10 or *R*10 and *R*4,*S*7,*R*9,*R*10 of 9H10epoZ are very unlikely to be correct. Thus, for 9H10epoZ four structures, with *R*4,*R*7,*S*9,*S*10, *R*4,*R*7,*S*9,*R*10, *R*4,*S*7,*S*9,*S*10 and *R*4,*S*7,*S*9,*R*10 configurations, remain possible.

To determine which configurations for 9HZ and 9H10epoZ are most likely correct, we performed protein homology modelling for ShZO. Systematic docking studies for both compounds (9HZ and 9H10epoZ) with all configurations tested for the CD spectra simulations (R4 stays fixed), including equatorial as well as axial orientation of the side chain with respect to the ring system, were carried out. The structure of the protein model used is shown in **Supplemental Figure 11**. The docking studies with all possible stereoisomers of zingiberene, show that the *R*4,*R*7 configuration is the only one adopting a docking pose with a short distance between a hydrogen atom bound to C9 and the reactive oxygen atom bound to the heme (Figure 5A). In all other configurations, docking arrangements with the cyclohexadiene ring moiety in proximity to the heme do not allow oxidation at C9. This is caused by the shape of the hydrophobic binding pocked formed mainly by L208, I116, V117, L362, and V363. In the case of the docked 4*R*,7*R*-zingiberene, the pro*S* H9 is at a distance of 2.5 Å, whereas the proR hydrogen is at a distance of 3.9 Å. Thus, the formation of 4*R*,7*R*,9*S*-hydroxy-zingiberene is highly favored. This conclusion is strongly supported by the results of the docking studies for 9-hydroxy-zingiberene with the 7*R*, 9R-, 7*S*, 9*S*-, and 7*S*, 9*R*-configurations. Only in the case of 4*R*,7*R*,9*S*-hydroxy-zingiberene does the docking pose (Figure 5B) allow an oxidation at the C10 atom exactly in the same orientation as that found for 4*R*,7*R*-zingiberene. Due to the orientation of the butene moiety, fixed by the restriction of accessible space caused by the spatially neighboring L208, the formation of 4*R*,7*R*,9*S*-hydroxy-10*S*,11-epoxy-zingiberene is the only possibility (Figure 5C).

**Figure 5.**
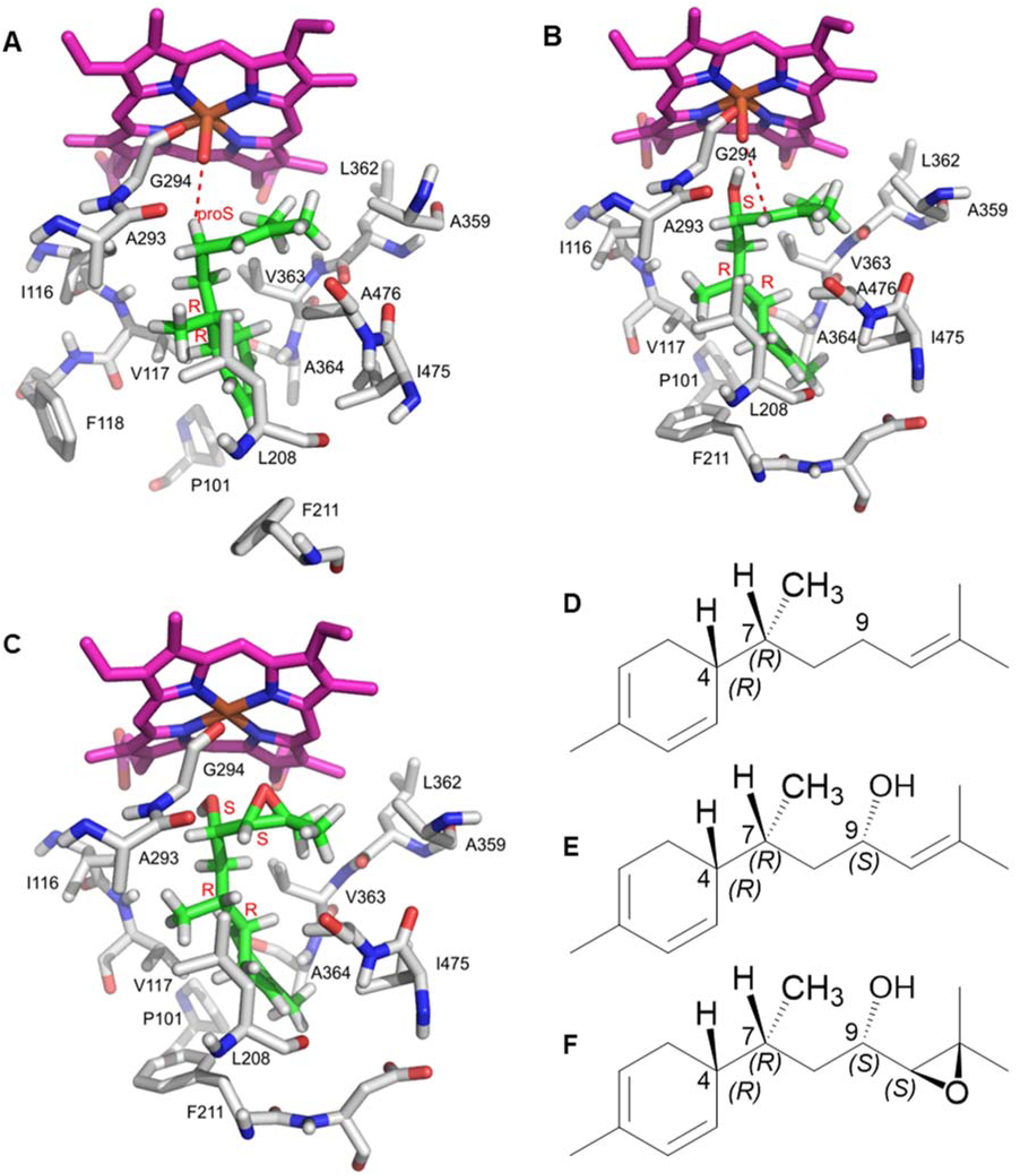
Determination of the absolute configuration of 9HZ and 9H10epoZ by molecular docking. The only possible docking arrangement of (*4R,7R)*-zingiberene to the active site of ShZO (ShCYP71D184) from *S. habrochaites* LA2167 is shown in **A** The pro*S* 9-hydrogen atom is in close distance (2.5 Å, red dotted line) to the reactive oxygen atom bound to the heme, favoring the formation of (*4R,7R,9S)*-9-hydroxy-zingiberene. The only possible docking pose of (*4R,7R,9S)*-9-hydroxy-zingiberene to the active site of ShZO is shown in B. It is in good agreement with the docking pose calculated independently for (*4R,7R)*-zingiberene. The reactive oxygen atom bound to heme is in close distance (3.4 Å) to the C10 carbon atom to favor the formation of tehepoxide with *S10* only. **C:** Formed (*4R,7R,9S,10S)-*9-hydroxy-10,11-epoxy-zingiberene in the active site. **D**: (*4R,7S)*-zingiberene. **E**: (*4R,7R,9S)*-9-hydroxyzingiberene. **F**: (*4R,7R,9S,10S)*-9-hydroxy-10,11-epoxy-zingiberene.

Thus, in summary and in agreement with the calculated CD-spectra, it is very likely that 4*R*,7*R*,9*S*-hydroxy-zingiberene and 4*R*,7*R*,9*S*-hydroxy-10*S*,11-epoxy-zingiberene are the enzymatically formed compounds (Figure 5D-F).

### 9-hydroxy-10,11-epoxyzingiberene exhibits toxicity against whiteflies

Next, we sought to evaluate the activity of these novel zingiberene derivatives towards whiteflies. To investigate the insecticidal properties of 7-*epi-*zingiberene and its derivatives, a series of no-choice whitefly bioassays were conducted using a realistic range of concentrations of the different fractionated compounds, applied to cultivated tomato leaf-discs. Concentrations were based on the quantities measured on the surface of *S. habrochaites* leaves (1-50 μg/cm^2^ for 7*epi*Z and 0.4-12 and 0.4-5 μg/cm^2^ for 9HZ and 9H10epoZ respectively). 48 Hours after exposure to individual treatments, the proportion of surviving whiteflies per cage was determined (**Supplemental Table 2**). Surprisingly, 7*epi*Z, the compound previously shown to be involved in whitefly resistance in *S. habrochaites* PI127826 (Bleeker et al., 2012), did not affect whitefly survival, even when applied at relatively high concentrations (Figure 6). In contrast, 9H10epoZ caused a significant and concentration-dependent reduction in whitefly survival (ANOVA, LDS post-hoc p < 0.01). This effect was however not found for 9HZ (Figure 6) and it appears therefore to be a very specific response.

**Figure 6:**
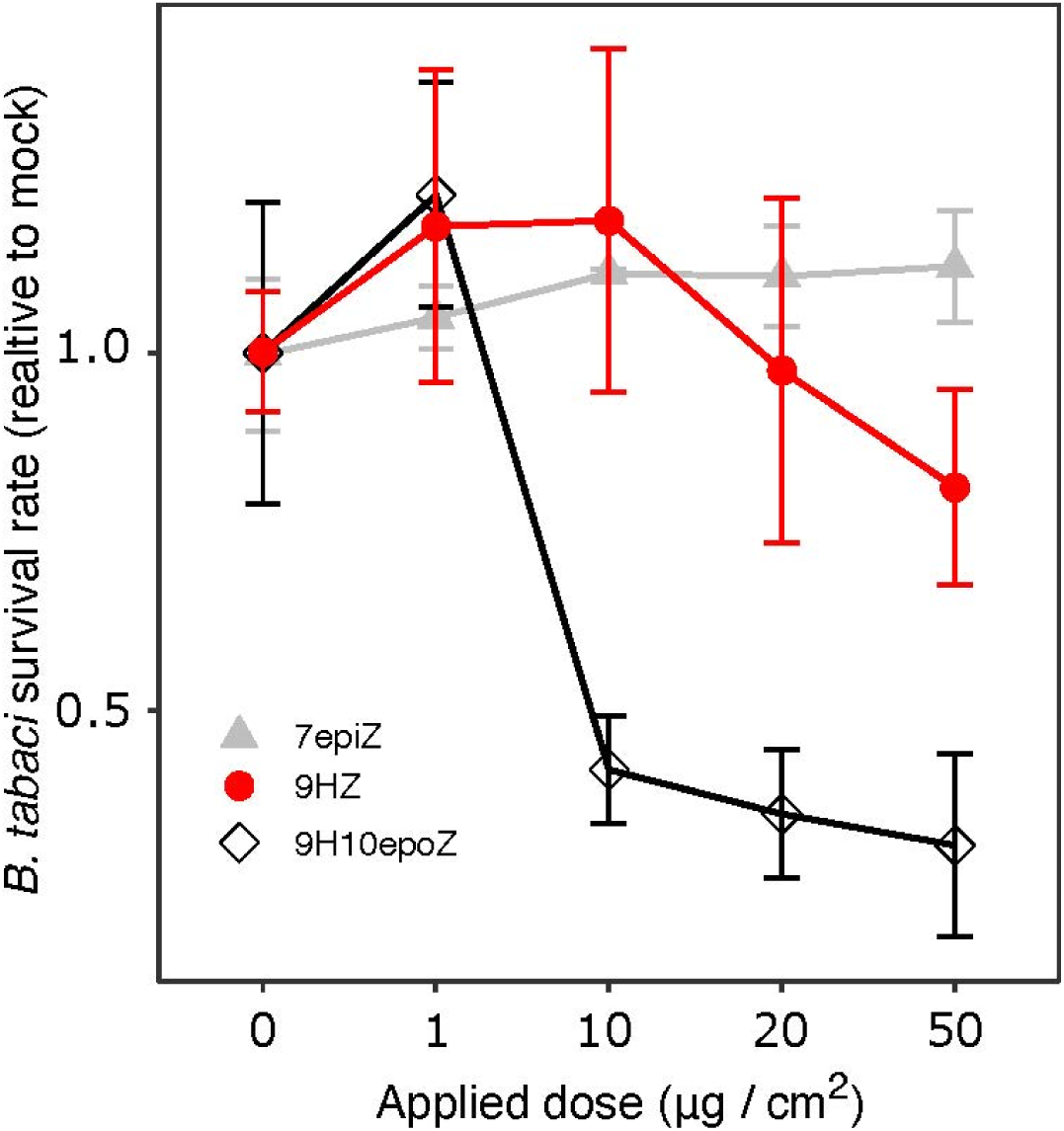
Whitefly toxicity assays of 4R,7R-zingiberene and its oxidized derivatives. Whitefly survival rates after 48 hours on treated leaf discs were normalised to the mean survival of the mock treatment (0 µg /cm^2^). Individual data points represent the average relative survival rate over 5 leaf discs ± SE.

### 7-*epi*-zingiberene and its derivatives have moderate antimicrobial activities

To evaluate the activity of 7*epi*Z, 9HZ and 9H10epoZ against microorganisms we performed growth inhibition assays with the bacterium *Bacillus subtilis*, the oomycete *Phytophthora infestans*, and the fungi *Botrytis cinerea* and *Zymoseptoria triticii*. All three compounds display growth inhibition activities against *B. subtilis*, with the strongest effect for 9H10epoZ and the weakest for 7*epi*Z (Figure 7A). Half-maximal response, or IC50 values, are 10.5 µM, 13.6 µM and 34.6 µM for 9H10epoZ, 9HZ and 7*epi*Z respectively. In contrast, in the assays with *P. infestans* only 9H10epoZ proves significantly active with an IC50 value of 55.8 µM, while 9HZ provides inhibition only at the highest concentration (200 µM) and 7*epi*Z has no inhibitory effect (Figure 7B). For *Z. triticii* a similar picture emerges, with 9H10epoZ being the most active compound with an IC50 of 33.0 µM (Figure 7C). Finally, against *Botrytis cinerea*, 9H10epoZ is again the most active compound with an IC50 of 21 µM, while 9HZ and 7-*epi*Z display an IC50 of 33.9 and 43.1 µM respectively.

**Figure 7:**
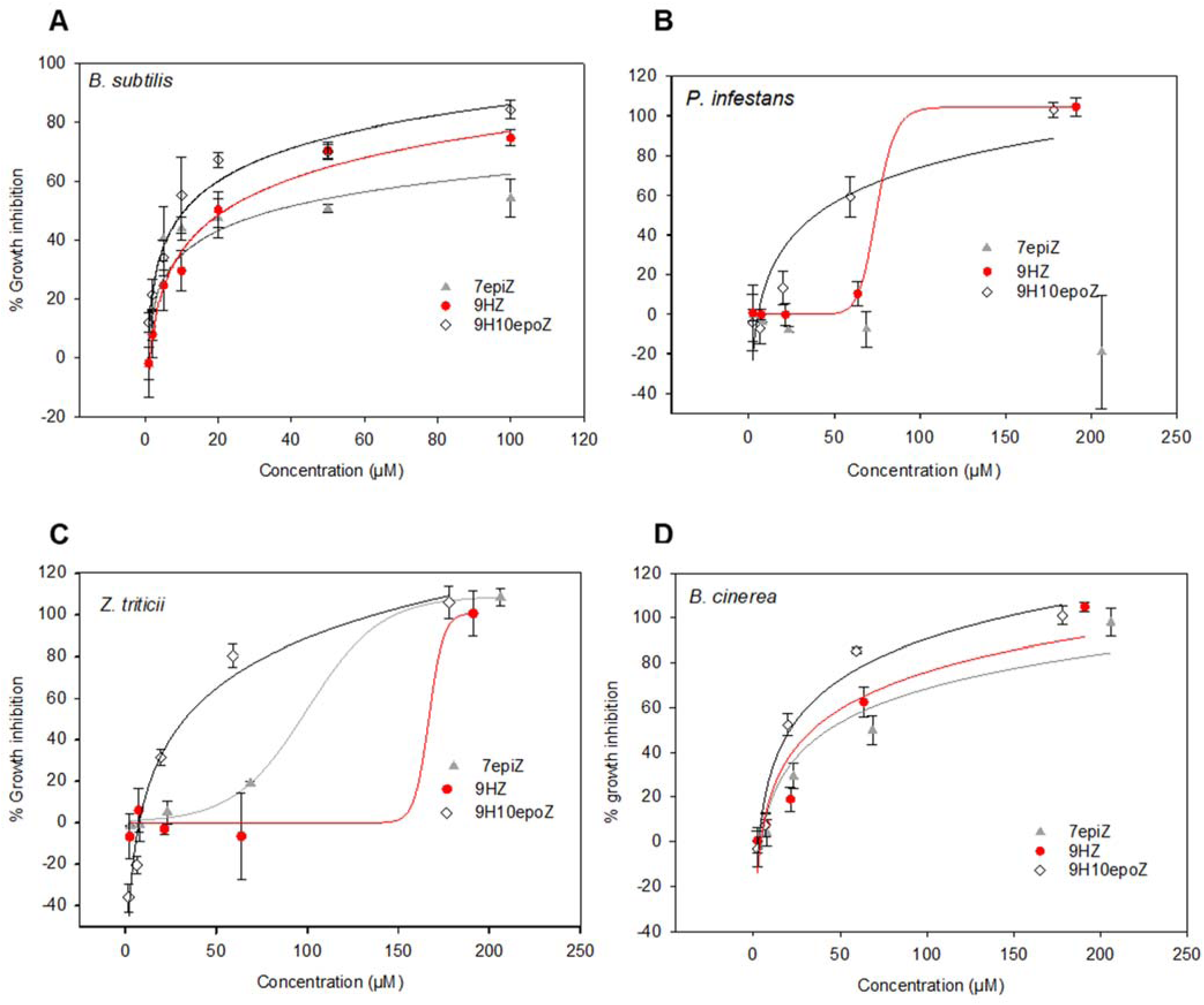
growth inhibition assays of 7epiZ, 9HZ and 9H10epiZ against various microrganisms. A: Assays against *Bacillus subtilis*. B: assays against *Phytophthora infestans*. C: assays against *Zymoseptoria triticii*. D. Assays against *Botrytis cinerea*.

### Discussion

Insect pests represent a significant threat for food security. Currently, in the majority of cases, they are dealt with by spraying synthetic insecticides, which are oftentimes not species-specific and therefore equally affect pests and beneficial insects, such as bees. This, and additional concerns regarding their impact on human health has led to the ban of some of these pesticides, such as the neo-nicotinoids, by environmental protection agencies in Europe and elsewhere (https://ec.europa.eu/food/plant/pesticides/approval_active_substances/approval_renewal/neonicotinoids_en). Furthermore, intensive use of these chemicals leads to the emergence of resistance in insect populations, rendering them ineffective. There is, therefore, a need for alternative strategies that limit the use of synthetic insecticides. One such an approach relies on the application of natural enemies of insects, such as parasitic wasps or pathogenic microorganisms (Lacey et al., 2015; Turlings and Erb, 2018). These strategies can work well with low levels of pest infestations but fail to cope with massive increase in populations under specific environmental circumstances. Another approach relies on the implementation of natural defenses of plants. In tomato, there is abundant literature highlighting the role of glandular trichome secretions in protecting plants against herbivore pests (Glas et al., 2012). However, these resistance traits are found exclusively in wild relatives of tomato. Breeding such metabolite traits into cultivated tomato is a challenging task because they are highly polygenic. First, specialized metabolites produced by plants are typically the products of pathways involving multiple enzymes. Although the underlying genes can be clustered in the genome as has been shown for a number of plant specialized metabolite pathways (Nutzmann et al., 2016; Nützmann et al., 2018), this is not always the case. Here, we show that the gene coding for zingiberene oxidase is on chromosome 1 in a different locus than the two genes required for the production of 7epiZ, which are clustered on chromosome 8. Secondly, as mentioned earlier, these metabolites are produced and stored in specialized tissues. Therefore, additional factors including trichome type, density and productivity need to be considered. There is strong support for a positive correlation between the level of resistance and the density of trichomes, which need to be of the correct type producing the compounds that are required for resistance. Tomato has four different types of glandular trichomes, with type I/IV producing acyl sugars and type VI terpenoids. The nature of the secretion by type VII trichomes is not clearly established, although analogous trichomes in tobacco produce antifungal peptides (Shepherd et al., 2005). Concerning type VI trichomes, a number of genes involved in their development have been identified (Schuurink and Tissier, 2020), but, to the best of our knowledge, genes controlling their density are not known yet. QTL analysis in crosses between *S. galapagense* and *S. lycopersicum* allowed the identification of a major locus on chromosome 2 contributing to both insect resistance and type IV trichome density together with the production of acyl sugars, but the genes still remain to be identified (Firdaus et al., 2013; Vosman et al., 2019). Yet another aspect is the capacity to produce and accumulate large quantities of specialized metabolites in trichomes. In some *S. habrochaites* accessions sesquiterpenoids produced by type VI trichomes can represent more 10% of the leaf dry weight (Frelichowski and Juvik, 2005), which is not explained only by the higher density of trichomes, but by the fact each trichome produces significantly more terpenes than in the cultivated tomato (Balcke et al., 2017). This could be due to the level of expression of the biosynthesis genes (Ben-Israel et al., 2009) but also to the capacity of type VI trichomes to store the metabolites in a cavity located between the glandular cells (Bergau et al., 2015; Bennewitz et al., 2018).

Here we identified two novel derivatives of 7-*epi*-zingiberene (7epiZ), 9-hydroxy-zingiberene (9HZ) and 9-hydroxy-10,11-epoxy-zingiberene (9H10epoZ) and the gene encoding ShCYP71D184, a P450 monooxygenase enzyme responsible for the biosynthesis of these two sesquiterpenoids in *S. habrochaites* LA2167. Although the absolute configuration of 9HZ and 9H10epoZ could not be determined directly, we inferred it by a combination of modeling the CD-spectra and docking 9HZ in the active site of ShCYP71D184. Based on these modeling approaches the only possible configurations were (4*R*,7*R*,9*S*)-7-*epi-*9-hydroxy-zingiberene and (4*R*,7*R*,9*S,10S*)-7-*epi*-9-hydroxy-10,11-epoxy-zingiberene. Direct demonstration of the absolute configuration of these compounds would require chemical synthesis or crystallization. In a recent publication Dawood and Snyder (2020) concluded that *S. habrochaites* contains 9HZ and 9H10epoZ based on spectral comparison with results that we released previously in a patent application (Dawood and Snyder, 2020).

Zingiberene-type sesquiterpenes have been identified in a number of plant and insect species. In ginger (*zingiber officinale*), α-zingiberene has a distinct configuration (4R, 7S) (Soffer and Burk, 1985) than 7-*epi*-zingiberene (*4R, 7R*) in *S. habrochaites* (Breeden and Coates, 1994). Ginger also produces *cis*- and *trans*-zingiberenol (also named bisabolenol) which are sesquiterpene alcohols with a hydroxyl group at position 1 instead of the double bond between C1 and C6 that is present in zingiberene (Khrimian et al., 2015). A mixture of stereoisomers of zingiberenols (*1RS,4RS,7S*) are sex pheromones produced by males of the rice stink bug (*Oebalus poecilus*) (de Oliveira et al., 2013) and interestingly two zingiberenol derivatives (*1S,4S,7R,10S and 1R,4S,7R,10S*) carrying an epoxide at position 10, like 9H10epoZ, are aggregation pheromones of the marmorated stink bug (*Halyomorpha halys*) (Khrimian et al., 2014). Marmorated stink bug is an emerging pest that is causing severe damage on a variety of fruit trees and horticultural crops (Haye and Weber, 2017). Although these pheromones have a different stereochemistry than the tomato zingiberene derivatives identified here, they support the importance of zingiberene-type compounds as insect pheromones. So far, no semiochemical has been identified for whiteflies (*Bemisia tabaci* and *Trialeurodes vaporariorum)* (Schlaeger et al., 2018) and it would be interesting to study the effect of the compounds identified here on the behavior of these important pests as well as other tomato herbivores.

The assays performed here point to a specific toxicity of one of the derivatives, namely 9H10epoZ, to which a clear, concentration dependent effect was observed against *B. tabaci* (Figure 6). The concentrations used were in line with the wild-tomato leaf surface. Thus, *S. habrochaites* LA2167 produces both a repellent (7epiZ) (Bleeker et al., 2011) and a compound that is toxic to whiteflies. Whereas it was previously assumed that wild tomato volatile sesquiterpenes inform the insect about the presence of lethal doses of the compound prior to landing {Bleeker, 2011 #5481}, here we show that 7epiZ, even at high concentrations, is not toxic to *B. tabaci*. Instead, the repellent volatile may signal the presence of a toxic derivative on the leaf surface. Previously we showed however that 7epiZ is toxic to spider mites using transgenic tomato lines that do not produce derivatives (Bleeker et al., 2012), further underlining the insect specificity of individual metabolites.

Furthermore, the 7epiZ derivatives and 9H10epoZ in particular, also showed moderate anti-microbial activity against several known plant micro-organisms including important pathogens like *B. cinerea* and *P. infestans* (Figure 7). *B. cinerea* and *P. infestans* are known tomato pathogens but *Z. triticii* is not. Thus, the anti-microbial activity of 9H10epoZ does not seem to be specifically directed against tomato pathogens. It remains debatable whether this anti-microbial activity is relevant in the field. In *S. habrochaites* LA1777 we previously estimated the amount of sesquiterpenes produced to be around 65 ng per type VI trichome (Balcke et al., 2017). Given that the head of type VI trichomes in *S. habrochaites* has a diameter of around 65 µm and that the volume of the storage cavity represents 65% of the trichome head (Bergau et al., 2015; Balcke et al., 2017), the average volume of the storage cavity is 93 465 µm^3^. Assuming that terpene production in LA2167 is equivalent to that of LA1777, this would correspond to a total sesquiterpene concentration of 3.1 M, or 1.03 M per sesquiterpene (7epiZ, 9HZ and 9H10epoZ, average MW= 220 Da) in the storage cavity, based on approximately equal concentration of the three major sesquiterpenes (see Figure 1). Thus, when the trichome head is ruptured, relatively high concentrations of these compounds are released, which could indeed exert inhibitory effects on pathogens present. Quantitative resistance to *P. infestans* was found in different accessions of *S. habrochaites* with QTLs located on chromosome 11 and 5 (Johnson et al., 2012; Haggard et al., 2013, 2014; Copati et al., 2019). These were done with LA1777 not with LA2167 however and further investigation on a putative contribution of sesquiterpenes of LA2167 to quantitative resistance to microbial pathogens should be carried out in the future.

In conclusion, we identified 7epiZ oxidation products and the enzyme responsible for their biosynthesis from an accession of the wild tomato *S. habrochaites.* This information can be used to breed tomato lines with improved performance against pests by reconstituting the whole pathway either by classical breeding, transgenesis or the CRISPR-Cas9-based knock-in technology (Dahan-Meir et al., 2018). Of these, the CRISPR-Cas9 approach is probably the most promising as it should lead to optimal gene expression levels and avoid the introgression of large chromosomal fragments around the genes of interest from *S. habrochaites*, which can lead to negative side-effects.

## Materials and Methods

### Plant Material and Growth

Seeds of *Solanum habrochaites* accessions LA2167, LA1777, LA1753, LA2650 and *S. lycopersicum* accession LA4024 were obtained from the Tomato Genetics Resource Center at UC Davis (http://tgrc.ucdavis.edu/). *S. habrochaites* accession PI 127826 was obtained from the USDA Germplasm (https://www.ars-grin.gov/). Plants were grown in a greenhouse with controlled conditions of 25°C and 55% humidity during the day, and 20°C and 75% humidity during the night. Plants were illuminated for 16 h (from 6:00 am to 10:00 pm) resulting in a light intensity of 5 to 25 Klux depending on weather conditions. The plants were watered once a week with a fertilizer solution (0.1% Kamasol Brilliant Blau, Compo Expert GmbH, Germany).

### Trichome Harvest

Leaves from 5-6 week old plants were removed from the plants and immediately brushed several times across the surface with a paint brush dipped in liquid N_2_ while keeping it right above a mortar containing liquid N2. The collection of trichomes from 20-30 leaves was sufficient to extract total RNA for cDNA synthesis. The harvested material was filtered through a steel sieve of 150 µm diameter (Atechnik, Leinburg, Germany) to further separate the isolated glandular trichomes from leaf debris and to eliminate non-glandular trichomes. After N_2_ evaporation, the collected trichomes were transferred with a cooled down spatula into a 1.5 mL reaction tube and stored at −80°C until further processing.

### RNA isolation and cDNA synthesis

RNA-isolation of leaf and trichome material from tomato plants was done using the RNeasy Mini Kit (Qiagen, Germany). The RNA concentration was measured with a Nanodrop spectrophotometer (Thermo Fisher Scientific) and the quality of the RNA was controlled with the QIAxcel Advanced system (Qiagen).

A DNase I digest was done using the Ambion DNA-free™ kit (Life Technologies – Carlsbad, USA) and 1 µg of total RNA and 1 µL of oligo-18dT primer was used for cDNA synthesis as follows. Nuclease-free water was added to a final reaction volume of 12 µL. Then 4 µL 5 × reaction buffer, 1 µL Ribolock RNase inhibitor, 2 µL deoxynucleotides (dNTPs) and 1 µL of RevertAid H Minus-MuLV reverse transcriptase (ThermoFisher Scientific) were added to a final volume of 20 µL. The mixture was then incubated at 42°C for 1 h and heated to 70°C for 10 min. The resulting cDNA was ready-to-use for PCR amplification reaction and for qRT-PCR expression analysis.

### Polymerase-chain reaction (PCR)

For DNA amplification from genomic DNA, cDNA or plasmid DNA for later cloning procedures different polymerase-chain reaction (PCR) methods were used. KOD Hot Start DNA Polymerase (KOD) (Sigma-Aldrich), Phusion high fidelity DNA polymerase (Phusion) (New England Biolabs) and the proofreading DreamTaq DNA-Polymerase (DreamTaq) (Thermo Fisher Scientific) were used following manufacturer’s recommendations. The following temperature program was applied including an initial denaturation step of 2 min at 95 °C and then 30-35 cycles of 30 s denaturation at 95°C, a 30 s annealing step at 55°C and elongation for 0.5 min/kb (KOD) or 1 min/kb (DreamTaq) gene length at 72°C with 5 min final extension step at 72°C. Oligonucleotide primers used for PCR amplification are listed in the Supplemental Data.

### Quantitative Real-Time PCR (qRT-PCR)

The *qRT*-PCR analysis was done with a Connect 96x Real-Time PCR system (BioRad). Input of each reaction was 3 μL of the cDNA reaction diluted 20 times. The reaction was done with the my-Budget 5x EvaGreen QPCR Mix II (Bio&Sell). The following program was applied including an initial denaturation step of 15 min at 95°C followed by 40 cycles with 10 s at 95°C and 20 s at 55°C. The melt curve was recorded after additional denaturing at 95°C for 30 s, cooling down to 65°C for 30s and heating up again to 95°C 0.1°C/s. The analysis was done with the Bio-Rad CFX manager software. For high-resolution melt (HRM) analysis gDNA (10 ng/μl) was used and the temperature program adjusted to 95°C for 15 min followed by 40 cycles with 10s at 95°C, 20s at 60°C and 72°C for 30s. In this case an extended melt curve was obtained by heating up to 95°C for 1 min, cooling down to 60 °C, 1 min hold at 60°C, followed by heating from 70 to 90°C at 0.01°C/s. The data was analyzed with the Precision Melt Analysis program (BioRad). The complete list of oligonucleotides used is provided in the Supplemental Data.

### cDNA Microarray hybridization

cDNA from leaves of the *S. habrochaites* accessions LA1753, LA2167, LA1777, LA2556 and LA2650 was hybridized to a custom designed microarray chip and the data analyzed as described in Balcke et al. (2017). Only the data for three genes are shown in Figure 3A.

### *S. habrochaites* LA2167 × *S. lycopersicum* LA4024 back-cross population and molecular mapping

To generate a back-cross population between accession LA2167 (*S. habrochaites*) and LA4024 (*S. lycopersicum*), pollen from the anthers of the stamen (male) of LA2167 was collected on a glass slide and used to pollinate emasculated flowers of LA4024. The procedure was repeated several times over 7 days to increase the chances of successful fertilization. Afterwards all flower parts from the cultivated tomato LA4024 except the pistil (female) were removed carefully and the collected pollen was transferred on to the stigma. This procedure was repeated several times with each pollinated flower over the following 7 days to ensure pollination. After fruit development all seeds were harvested and sowed after 1 week of drying at room temperature. The progeny was then analyzed by GC-MS and HRM analysis genotyping for at least 8 independent loci. Only plants that had zingiberene and its derivatives and were heterozygous for the markers were kept for the back-crosses. After flowering pollen of the F1 plants was collected and used to pollinate the cultivated tomato. Seeds were harvested and each BC1F1 plant was then individually characterized genotyped and analyzed by GC-MS as for the F1 plants. Plants that had the phenotype and genotype of LA4024, i.e. resulting from self-fertilization of LA4024, were discarded.

### Molecular mapping

114 markers were used. The complete list is provided in **Supplemental Table 2**. 25 were High Resolution Melt (HRM) markers, the others were iPLEX™ assay markers. The iPLEX™ assays were performed at ATLAS-Biolabs (Germany). The High Resolution Melt assays were performed as described in Bennewitz et al. (2018).

### Leaf surface extracts and analysis by GC-MS

Surface extracts were collected from 6 leaflets of circa 5 cm length by adding 2 ml n-hexane and shaking for 1 minute. The hexane was taken off and centrifuged at 16,000 × g for 90 seconds to remove debris. 1 µl of the hexane supernatant was then injected in a Trace GC Ultra gas chromatograph coupled to an ISQ mass spectrometer (Thermo Scientific). GC-MS analyses were performed as described in Brückner et al. (Brückner et al., 2014).

### Purification of 9HZ and 9H10epoZ

To get enough soluble material for normal solid phase extraction (SPE) chromatography and further experiments (bioassays and NMR analysis) around 500-1000 leaves of the *S. habrochaites* accession LA2167 were extracted with 500 mL of n-hexane for 5 min. The organic solvent was then gently removed by a rotary evaporation system Rotavapor R-114 (Fluke, Germany) at room temperature. The dried sample was then resolved in 50 mL n-hexane and stored at −20°C. In parallel the SPE column was packed with 15 mg CHROMABOND silica adsorbent material (Machery-Nagel) and equilibrated several times with 50 mL n-hexane. After preparation the sample was applied on to the column. Afterwards the column was washed six times with an 90:10:1 mixture of n-hexane:ethyl acetate:methanol. A second elution step was then carried out with a 50:50:1 mixture of n-hexane:ethyl acetate:methanol. 50 mL fractions were collected and cooled down at 4°C until further analysis.

### Nuclear Magnetic Resonance Analysis

NMR spectra were recorded on an Agilent (Varian) VNMRS 600 NMR spectrometer at 599.832 MHz (^1^H, ^2^D) and 150.826 MHz (^13^C) using a 5 mm inverse detection cryoprobe. The samples were dissolved in C_6_D_6_. Chemical shifts were referenced to internal TMS (δ ^1^H 0 ppm) and internal C_6_D_6_ (^13^C 128.0 ppm).

### Circular Dichroism

CD spectra were recorded on a JASCO-J815 CD spectrometer using n-hexane as solvent.

### Molecular Cloning

*Golden Gate* cloning was used for the preparation of all plasmid constructs reported here (Engler et al., 2008; Weber et al., 2011; Werner et al., 2012; Engler et al., 2014). The complete list of constructs used here is provided in the Supplemental Data. The zFPS coding sequence was codon optimized (see Supplemental Data) for *N. benthamiana*, while ShZS (Zingibernee synthase) and ShCYP71D184 were amplified as cDNA from LA2167.

### Transient expression in *Nicotiana benthamiana*

T-DNA constructs containing cDNAs for zFPS, ShZS, ShZO (CYP71D184) and the Arabidopsis cytochrome P450 reductase (ATR1) all under the control of the 35S promoter (see list of contructs in the **Supplemental Data**), were transformed into *Agrobacterium tumefaciens* strain GV3101 (pMP90) by electroporation. *A. tumefaciens* strains containing the respective T-DNA were grown in a 5 mL LB medium pre-culture for 24 h at 28°C under constant shaking at 180 rpm. 0.5 mL of this pre-culture was then used for inoculation of 50 mL LB medium and grown for 16 h at 28°C. The *A. tumefaciens* cells were then mixed up in different combinations of expression constructs and harvested by centrifugation for 30 min at 1800 g at 4°C at a global OD_600_ of 0.6. In combinations with fewer genes, the mixture was complemented by the addition of an equivalent amount of an *A. tumefaciens* strain expressing GFP with the 35S promoter. The cells were resuspended in infiltration buffer (5% sucrose (w/v) and 4.3 g/L Murashige and Skoog basal salt mixture) containing 20 µM acetosyringone. Leaves of 4-5-week old *N. benthamiana* plants were infiltrated by pressing a 1 mL syringe onto the abaxial side of the leaves followed by slowly injecting the *A. tumefaciens* suspension. After complete infiltration of 3-4 leaves the plants were brought back into growth chambers for additional 5 days after which they were extracted for GC-MS analysis.

### Computational methods for the calculation of CD spectra

Conformational analysis of all compounds under investigation was performed using MMFF94 molecular mechanics force field (Halgren, 1999) and low-mode molecular dynamics simulations within the Molecular Operating Environment (MOE) software (v2015.10; Chemical Computing Group Inc.: Montreal, QC, Canada, 2015). The force field minimum energy structures were subsequently optimized by applying the density functional theory (DFT) using the BP86 functional with the def2-TZVPP basis set (Perdew, 1986; Becke, 1988; Schäfer et al., 1992; Weigend and Ahlrichs, 2005; Karton et al., 2008) implemented in the ab initio ORCA 3.0.3 program package (Neese, 2012). The influence of the solvent n-hexane was included in the DFT calculations using the COSMO model (Sinnecker et al., 2006) to include any influence of the solvent. The quantum chemical simulation of the UV and CD spectra was also carried out using ORCA. Therefore, the first 50 excited states of each enantiomer and conformation were calculated by applying the long-range corrected hybrid functional TD CAM-B3LYP9 with the def2-TZVP(-f) and def2-TZVP/J basis sets (Perdew, 1986; Becke, 1988; Schäfer et al., 1992; Weigend and Ahlrichs, 2005; Karton et al., 2008). The CD curves were visualized with the help of the software SpecDis 1.64 (https://specdis-software.jimdofree.com/) (Bruhn et al., 2013) from the calculated rotatory strength values using a Gaussian distribution function at a half-bandwidth of σ = 0.3 eV. The single spectra of the individual conformers were summed up according to their contribution to Boltzmann-statistical weighting (as derived from the single-point energy calculations), wavelength shifted, and compared with the experimental spectra.

### Protein Homology Modelling

Protein homology modelling of ShCYP71D184 was automatically performed with YASARA (Krieger et al., 2009). After search for template proteins in the protein database (Berman et al., 2000), 100 models were created based on alternative sequence alignments including secondary structure predictions and comparisons with found appropriate X-ray protein structures. The model based on the x-ray structure of 3CZH (CYP2R1) appeared as best model with a sequence identity of 28% and a sequence similarity of 44.9%. A final model was formed by merging short loop sequences from several other proteins. The quality of the model was checked for native folding by energy calculations with PROSA II (Sippl, 1990, 1993) and for stereo-chemical quality by PROCKECK (Laskowski et al., 1993). The Ramachandran plot showed 88.2 of all amino acid residues in most favored areas and five outliers located in loops outside the active site. All other stereo-chemical parameters are in allowed regions for good quality. The z-score value of −8.8 for combined potential pairs resulting from the PROSA analysis indicates a native like fold. In the final model, the co-substrate heme was automatically overtaken from the X-ray structure and was inserted in the model. Manually an oxygen atom was added to the in heme coordinated iron ion using the “molecular operating environment” program package MOE 2015.1001 (https://www.chemcomp.com/).

Energy optimized structures of zingiberene all four alternative configurations (except fixed R-configuration for C4, 7R and 7S zingiberene; 7R, 9R-, 7R, 9S-, 7S, 9R-, and 7S-9S-hydroxy-zingiberene) including for each an equatorial as well as an axial orientation of the side chain for both compounds were used for docking studies. Docking was performed with GOLD and chemscore scoring function allowing for each structure the output of 30 poses. For all optional parameters standard settings were applied. All these produced poses were analyzed by measuring the distances between the oxygen atom bound to iron and both hydrogen atoms at C9 (proS, and proR) in case of zingiberene and for hydroxy-zingiberene the distances to C10 and C11.

### *Bemisia tabaci* toxicity assay

SPE purified fractions were dried under N2 and dissolved in 1.7 mM TritonX-100 followed by thorough mixing using a vortex. 20 μL of diluted fraction, containing the appropriate concentration of terpenoids, was applied to the abaxial side of freshly harvested leaf discs (1.5 cm Ø) from five-week-old *S. lycopersicum* (cv. C32) plants. Leaf discs treated with 0 to 50 μg terpenoids per cm^2^ (n = 5) were air dried for 30 minutes prior to starting the experiment. 15 whiteflies (*B. tabaci*) were collected from a cucumber rearing and placed in small containers (Greiner analyzer cups; 2.9 cm Ø × 5.2 cm) with a Ø 1 cm opening in the lid. The opening was covered with a leaf disc facing the treated side inside of the cup. The leaf disc was subsequently covered with moist filter paper (Whatman) and sealed with Parafilm®. Cages were placed for 48 hours in a climate chamber at 24°C where after dead and alive whiteflies were counted under a stereomicroscope. The survival rate per cage was determined and, by means of standardizing separate experiments, survival rates were calculated relatively to the mean survival of the mock controls.

### Antibacterial Assay

To determine the antibacterial activity of 7epiZ and its derivatives, this assay was done with a genetically modified version of the gram-positive *Bacillus subtilis* strain 168 expressing the yellow-fluorescent protein YFP (Veening et al., 2004).This protein is under the control of the abrB promotor, which is only active during the exponential growth phase of *Bacillus subtilis*, which allows determining the exact number of bacteria and their vitality by direct fluorescence measurements. In this case the fluorescence could be used indirectly as a “benchmark” for the growth inhibition induced by the test compounds (Michels et al., 2015). The *Bacillus subtilis* 168 strain was grown on TY media with 1% tryptone, 0.5% yeast extract, 1% NaCl and chloramphenicol (5 ug/ml) for selection. After overnight growth on solid media (1.5% agar) (Fluka) at 30 °C, pre-cultures (50 ml) for the antibacterial assay were inoculated from plate and incubated for 24 h at 30°C in liquid media without shaking. The cell density was adjusted to 1.6 × 10^5^ cells/ml resuspended afterwards in 10 mL fresh TY media and used for the bacterial assay. The assay was done in black flat-bottom 96-well microtiter plates (BD Falcon). In addition to a negative control (300 µL TY media) four different controls were carried along with each plate. First the auto-fluorescence of the single compounds (K1, n = 1) was measured; then the bacterial suspension alone (K2, n = 6); the reference control was a mixture of pure methanol and inoculum (K3, n = 6) equivalent to 100% growth; Finally, K4 was a positive control containing erythromycin, a common antibiotic, which was dissolved in pure methanol. The samples themselves were a mixture of the tested compound at different concentrations (1 to 100 µM) and the bacterial inoculum. The fluorescent measurement was done with a microtiter plate reader Tecan GENios Pro (λ_ex_ = 510 nm, λ_em_ = 535 nm, bandwidth=10 nm, gain 45, 60, 10 flashes, integration time 40 μs, 50% mirror, 3 × 3 quadratic measurements per well, temperature 27 °C, shaking 10 s low intensity, settle time 1 s) at the time point t = 0 (t_0_) after inoculation with *Bacillus subtilis* and again under the same specifications after 15 h (t_15_) incubation at 30 °C. Further calculations of the percentage of growth inhibition relative to K3 were done referring to the following equation 1: growth inhibition [%] = (1-100x (x_sample_ [t_15_-t_0_]/x_k3_ [t_15_-t_0_])) × 100

### Assays for Growth Inhibition of Plant Pathogens

The strains used were *Botrytis cinerea* Pers., *Septoria tritici* Desm., and *Phytophthora infestans* Mont. and the assays were performed in 96-well microtiter plate assay. The experimental setup was done according the fungicide resistance action committee FRAC (www.frac.info/). Except some minor modifications in the protocol. 7-epi-zingiberene, 9-hydroxy-zingiberene and 9-hydroxy-10,11-epoxy-zingiberene were tested in a dilution series of a stock solution from 0.52 µg/ml to 42 µg/mL t. The compounds were diluted in DMSO with a final concentration of 2.5% in the assay. Additionally, DMSO was also used as negative control, whereas pyraclostrobin (Sigma-Aldrich), which is a commercially available fungicide, was used as positive control. For each concentration three biological replicates were measured. Pathogenic growth was determined by measuring the optical density (OD) at λ = 405 nm 7 days after inoculation with a microtiter plate reader GENios Pro (Tecan) doing 5 measurements per well using multiple reads with 3 × 3 quadratic measurements per well).

## Acknowledgments

We would like to thank the IPB for funding this project and the greenhouse personnel of the IPB (Petra Jansen, Philip Plato and Sabine Voigt) for taking care of the plants.

## Author contributions

The project was conceived and supervised by AT. SZ performed the identification and purification of the zingiberene derivatives, generated the backcross population, analyzed the BC1F1 plants by GC-MS and carried out the molecular mapping and all the cloning and the transient expression in *N. benthamiana*. WB did the CD-spectrum simulation and the 3D-protein modeling and docking. AP performed the NMR analysis. BA provided support for the design of molecular markers. RK and PB performed the whitefly assays. AT wrote the manuscript, which all authors revised and approved.

